# Human striatal population state dynamics

**DOI:** 10.64898/2026.06.17.733040

**Authors:** Cole Korponay, Elliot A Stein, Thomas J Ross, Amy C Janes

## Abstract

Animal models reveal that striatal projection neurons (SPNs) fluctuate between discrete electrophysiological states with distinct levels of cortical interaction. These dynamics and their behavioral relevance remain uncharacterized in humans. Leveraging neurobiologically informed modeling of over 3 billion voxel-frame-wise striatal coactivation profiles with cortex in functional magnetic resonance imaging (fMRI), we identified population-level striatal states in humans resembling canonical SPN states that reorganized systematically with task demands, arousal and behavior. A background of low- and moderate-coactivation “down-like” and “up-like” striatal rest states with high transition reciprocity modulated task reaction times and reward reactivity. Meanwhile, sparse, disproportionately high-magnitude ‘bursts’ of striatal coactivation with cortex, which emerged preferentially from the up-like rest state and whose cortical input composition varied with task context, tracked task engagement and arousal level. Findings bring a critical feature of corticostriatal neurobiology into systems-level view in humans and reveal a subthreshold state architecture whose balance encodes behaviorally relevant information.

## Introduction

The striatum and frontal cortex work in concert to coordinate a wide range of adaptive functions in the motor, cognitive and limbic domains^1^. Facilitating the interaction of these structures is a vast network of cortical projections that monosynaptically innervate the striatum in a topographic manner^1–3^. Individual striatal territories receive appreciable projections from a distributed set of cortical regions^3–6^; these projections and their relative strengths comprise a striatal territory’s cortical “connectivity profile”^1,7–10^. These projection profiles define the cortical input channels through which distinct striatal territories integrate information relevant to action selection, reward valuation, cognitive control, and sensorimotor execution, providing an anatomical scaffold for functional specialization.

Signal transmission across corticostriatal projections can be inferred from correlated fMRI blood oxygen level dependent (BOLD) signal activity (i.e., time-averaged functional connectivity, FC) between anatomically connected regions of cortex and striatum^11,12^. Though FC is a non-directional measure, and striatal activity also indirectly influences cortical activity via cortico-basal ganglia-thalamo-cortical loops^1^, the FC strength of cortex-striatum node-pairs is strongly correlated with the strength of their direct structural connections^12–16^. Thus, a topographic map of corticostriatal FC largely mirrors that of the underlying structural connectivity map^12,15,16^. In aggregate, these corresponding static maps of corticostriatal circuits frame much of the current understanding of interactions between these structures at the systems level in humans.

Yet, while the gross structural projections underlying corticostriatal circuitry have a stable architecture, several lines of evidence suggest that striatal connectivity profiles with cortex are functionally dynamic and flexible. First, recent advances in time-resolved fMRI connectivity analysis have revealed that interregional functional connection strengths fluctuate substantially across time, rather than remaining stable around a fixed mean^17–19^. Second, animal electrophysiological models have long shown that striatal projection neurons (SPNs, the striatum’s primary recipient of cortical input, and comprising ~95% of all striatal neurons^20^) fluctuate between three energetic states that exhibit distinct levels of engagement with frontal cortex. SPNs spend most of the time toggling between two resting states: a hyperpolarized “down” state characterized by negligible interaction with and responsivity to cortex, and a depolarized “up” state characterized by moderate interaction with cortex and appreciable responsivity to marginal excitatory input^21,22^. These states have similar per entry dwell times (~300-400ms) and regularly fluctuate between one another^23^. The third SPN state is an active state, wherein an SPN in the “up” rest state receives sufficient marginal excitatory input to fire action potentials^24,25^. This multi-state system facilitates sparse temporal coding that allows for precise and flexible behavior.

Despite their centrality to brain function, striatal state dynamics remain essentially unmodeled and unexplored in humans, precluding study of their role in human behavior and disease. While the fMRI blood oxygen level dependent (BOLD) signal is not a direct readout of neuronal voltage levels^26^, emerging high-temporal resolution methods for tracking interregional BOLD coactivation magnitudes^27^ may allow for indirect discrimination of striatal activation states (at the population/mesoscale/voxel-wise level) via their characteristic differences in cortical coactivation levels. We tested this supposition to identify fMRI-based signatures of striatal population state dynamics in humans, track and characterize their features across resting-state and task conditions, and evaluate their links with arousal and individual differences in several behavioral phenotypes.

To do so, we leveraged the established anatomical and electrophysiological features of corticostriatal circuitry, together with recent advances in fMRI signal processing^28^ and neurocircuit modeling^7,27^ to infer voxel-scale striatal population state dynamics from cortical coactivation profiles sampled every 0.72 seconds across the full striatum over 65 total minutes of scanning in six acquisitions. The spatial resolution (2mm isotropic) and temporal blurring of the fMRI hemodynamic response function^29,30^ and denoising processes yield a measurement unit (i.e., voxel-frame) comprising seconds-worth of neuronal activity averaged across several hundred thousand neurons. We found that even at this mesoscale, the dynamics of striatal voxel coactivation profiles with frontal cortex recapitulate many of the features of canonical striatal electrophysiological states and reorganize systematically across arousal levels and task context in ways that track individual differences in behavior. This identification of neurobiologically and behaviorally meaningful substructure in striatal fMRI signal provides a foundation for probing the roles of striatal population state dynamics in human behavior and disease.

## Results

The goals of this investigation relied on quantification of within-scan temporal dynamics of BOLD-based striatal-cortical interregional coactivation. As such, fMRI data were denoised using a combination of nuisance regression and regressor interpolation at progressive time delays (RIPTiDe)^31^, a pipeline that our recent work demonstrated removes the greatest extent of temporally-structured noise (i.e., achieves maximal within-scan reliability of the BOLD signal) among widely used fMRI denoising strategies^28^.

### Modeling and Identifying Structure in Corticostriatal Coactivation Dynamics

Retrograde anatomical tract-tracing data in non-human primates show that while a given unit of striatal space can receive non-negligible input from a dozen or more frontal cortical regions, a majority of input usually comes from the five strongest regions^4,32^. As such, we first operationalized each striatal voxel’s connectivity profile as its set of FC magnitudes with the five ipsilateral frontal cortical regions of interest (ROIs) with which it has the strongest time-averaged FC with (**Fig. 1a**), using the Schaefer 100-parcel, 7-network atlas^33^ (**Fig. S1**). For computational tractability, we focused only on the right hemisphere. For each right striatal voxel, we computed its framewise coactivation strength^27^ with each of the five cortical ROIs in its connectivity profile, using the element-wise product of the striatal voxel’s z-scored BOLD timeseries and the cortical ROI’s z-scored BOLD timeseries (**Fig. 1b**). This produced a five-dimensional coactivation strength vector for each striatal voxel at each scanning frame/TR (**Fig. 1c**). Using this framework, we computed the framewise coactivation profiles (measured every 0.72 seconds) of 1710 striatal voxels across 1180 timepoints in 407 quality-controlled subjects who had complete data from four 15-minute resting-state fMRI acquisitions in the Human Connectome Project-Young Adult cohort, yielding 3,284,978,400 framewise, voxel-wise striatal coactivation profiles. These coactivation profiles were subsequently clustered to identify recurring states (**Fig. 1d**).

**Figure 1.**
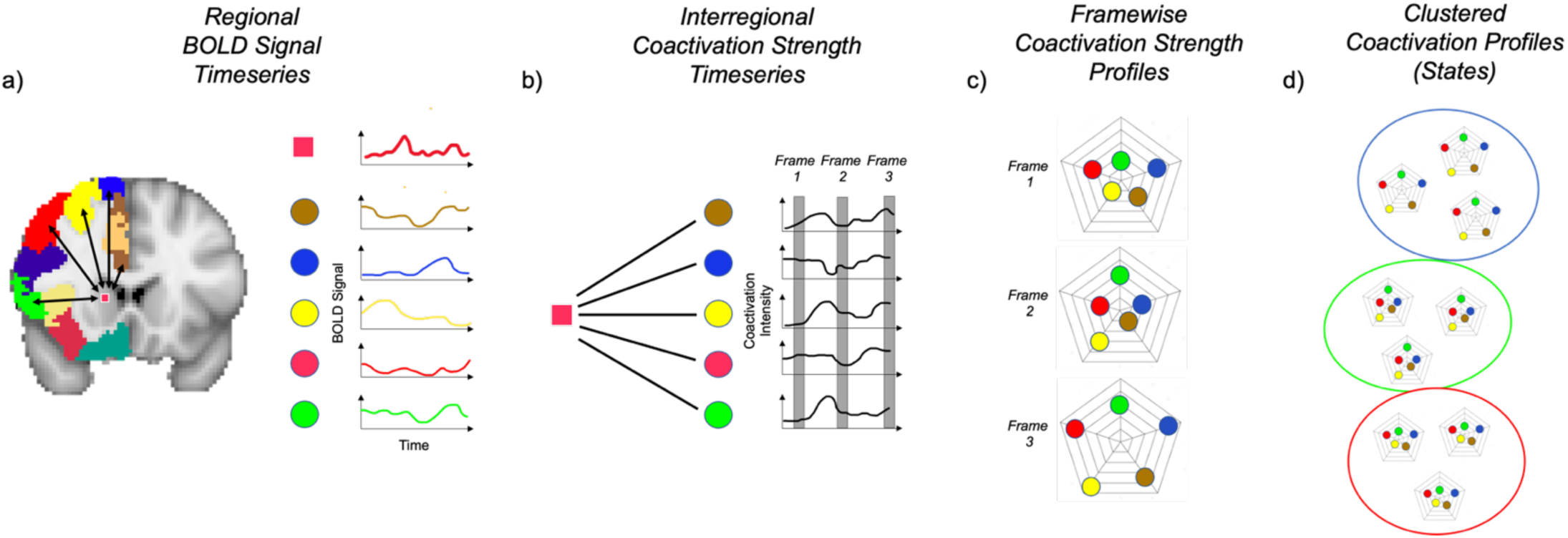
Identifying voxel-wise striatal coactivation states. a) The five strongest frontal cortical inputs to each striatal voxel were identified using static FC. The BOLD timeseries of a striatal voxel and its five strongest cortical inputs were used to compute b) five coactivation timeseries, encoding the coactivation magnitude of each cortico-striatal node-pair at each time frame. Each temporal slice through the five timeseries represents c) the “coactivation profile” of the striatal voxel at each point in time. d) Clustering was applied to these coactivation profiles across all time points, striatal voxels and subjects to identify recurring states.

For clustering, we employed a two-step procedure. Electrophysiological data demonstrates that striatal projection neurons spend most of their time in one of two non-firing “rest” states (i.e., down and up states) and only rarely enter a firing state^21^. At the same time, prior fMRI work characterizing framewise interregional coactivation has observed a subset of timepoints with disproportionately high-amplitude coactivations (i.e., coactivation “bursts”) that coincide with frames of high BOLD signal amplitude and carry high functional relevance^19,34^. In the case of structures with known monosynaptic connectivity, coactivation bursts between two such structures could signify periods when an activated upstream structure (e.g., cortex) is contributing to activation of a downstream structure (e.g. striatum). The properties of these coactivation bursts (i.e., rare in frequency and disproportionate in magnitude) are not well-suited to identification by common data-driven clustering algorithms applied to fMRI data (e.g., centroid-based or modularity maximization clustering)^35,36^. Indeed, model comparison analyses demonstrated that the distribution of coactivation profile strengths was substantially better fit by a two component Gaussian mixture model (GMM; Bayesian Information Criterion (BIC) = 97534) with distinct regimes (i.e., rare, disproportionately high-amplitude coactivation bursts and frequent, low-amplitude coactivation “rest” states) than by a one component GMM (BIC = 279800) representing a single continuous distribution. Moreover, null modeling (see Methods) indicated that the empirical distribution departed from unimodality more strongly than expected from autocorrelated noise alone (empirical ΔBIC(2-1) = −182,266; mean null ΔBIC(2-1) = −116,951 ± 851.8).

As such, we first applied a two-component GMM to the coactivation profiles to separate disproportionately high-coactivation “burst” states from low-coactivation “rest” states. We then applied unsupervised Louvain modularity maximization^37,38^ to the remaining low-coactivation frames to identify rest state subtypes (described in detail in Methods). To establish a standardized reference scale across rest and task acquisitions, clustering of coactivation profiles was performed on a reference resting-state acquisition, and the resulting state centroids and burst thresholds were then applied to classify coactivation profiles in the remaining three resting-state acquisitions and two Gambling-task acquisitions.

### Striatal State Dynamics at Rest

This analysis revealed that, across the four resting-state fMRI acquisitions, human striatal voxels spend a mean (± SEM) of 14.18% ± 0.05% of the time in a high-coactivation “burst” state and 85.82 ± 0.19% of the time in one of three low-coactivation “rest” states (**Fig. 2, Supplementary Videos 1-2**). By comparison, refitting the two-component GMM to the surrogate null data yielded a coactivation burst state prevalence of 39.01% ± 0.59%. This marked discrepancy (39.01% null versus 14.18% empirical) indicates that the empirical burst state threshold is not a trivial consequence of the GMM procedure or temporal autocorrelation alone but reflects distributional structure specific to empirical corticostriatal coactivation.

**Figure 2.**
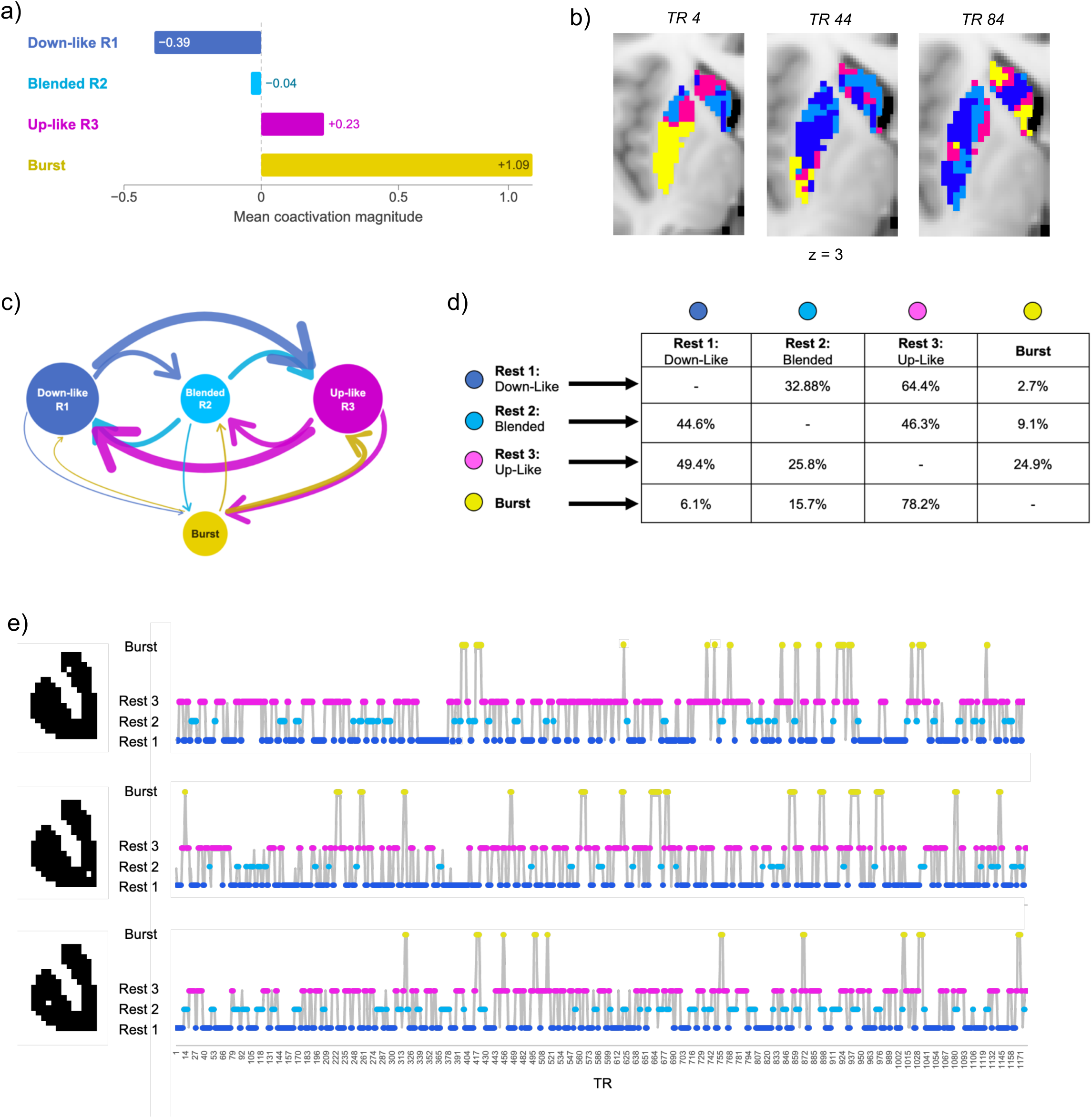
Quantitative profiling of striatal coactivation states. a) The mean coactivation magnitude (*z*-score) of striatal voxels with their five strongest cortical inputs – averaged across all striatal voxels, timepoints and subjects – for each of the four identified striatal coactivation states. b) Axial views of the striatum, at frames separated by 40 TRs, in a randomly selected subject illustrating the spatial structure and distribution of coactivation states over time. c) Out-of-state transition structure; lines weighted by count of out-of-state transitions and node size weighted by occupancy rate during an average resting-state scan. d) Probabilities of each out-of-state transition. e) For randomly selected voxels in the caudate, putamen and nucleus accumbens in a randomly selected subject, the temporal dynamics of each voxel’s coactivation state across a 15-minute resting-state fMRI scan.

In the coactivation burst state, a striatal voxel’s BOLD signal shows disproportionately high coactivation with that of its frontal cortical inputs (mean coactivation magnitude = 1.09 ± 0.77) (**Fig. 2a**). In rest state 1 (occupancy likelihood: 35.78 ± 0.08%) a striatal voxel’s BOLD signal shows minimal or negative coactivation with that of all or most of its five strongest frontal cortical inputs (mean (± SD) coactivation magnitude = −0.39 ± 0.47) (**Fig. 2a**). In rest state 2 (occupancy likelihood: 17.24 ± 0.07%), a striatal voxel’s BOLD signal shows moderate coactivation with that of some of its five inputs and minimal or negative coactivation with that of other inputs (mean coactivation magnitude = −0.04 ± 0.23). In rest state 3, (occupancy likelihood: 32.80 ± 0.04%), a striatal voxel’s BOLD signal shows moderate coactivation with that of all or most of its five inputs (mean coactivation magnitude = 0.23 ± 0.19). As such, we labeled rest states 1, 2 and 3 as “down-like”, “blended”, and “up-like”, respectively. The distributions of mean coactivation magnitudes for each state had distinct peaks but also some overlap (**Fig. S2**). Mean dwell times were 5.54 ± 0.01 TRs (3.99 sec) for down-like rest state 1, 3.46 ± 0.01 TRs (2.49 sec) for blended rest state 2, 3.88 ± 0.01 TRs (2.79 sec) for up-like rest state 3, and 5.23 ± 0.01 TRs (3.77 sec) for the burst state. States formed spatial clusters whose structure fluidly reconfigured over time (**Fig. 2b**).

All four states functioned as local attractors with high self-transition probabilities: P(rest 1→rest 1) = 0.820, P(rest 2→rest 2) = 0.712, P(rest 3→rest 3) = 0.743, P(burst→burst) = 0.809. When transitioning out of state (**Figs. 2c-d**), rest 1 and rest 3 showed strong bidirectional coupling: 64.4% of out-of-state transitions from rest 1 led to rest 3, and 49.4% of out-of-state transitions from rest 3 led to rest 1. Out-of-state transitions into burst were rare overall (2.7%, 9.1%, and 24.9% from rest 1, rest 2, and rest 3, respectively), but 68% of all rest-to-burst transitions originated from rest 3, consistent with bursts representing transient amplifications of the up-like configuration. The temporal dynamics of these striatal voxel states over the course of a full 15-minute resting-state scan are visualized for randomly selected voxels in **Fig. 2e**.

Reliability of the striatal state dynamics properties (e.g., occupancy rates, transition probabilities, dwell times) across repeated scanning acquisitions was quantified using intraclass correlation coefficients (ICCs). For each property we computed single-run ICC(A,1) (two-way random, absolute agreement, single measure) and ICC(A,k) (reliability of the *k*-run average) across the four resting-state runs. Single-run reliability ICC(A,1) ranged from 0.17 to 0.36, with transition probabilities and dwell times having higher reliability than occupancy rates. Averaging across the four rest runs substantially improved reliability, with ICC(A,k) rising to 0.64–0.69 for transition probabilities and dwell times and to 0.51–0.58 for occupancy rates.

### Coupling of Striatal State Dynamics to Arousal Level and Task Performance

Recent work demonstrates strong links between resting-state functional connectivity dynamics and arousal level^39^ even after physiological denoising^40^, implicating genuine neuronal dynamics in arousal-related BOLD signal variation. As such, we used linear mixed-effects modeling to disentangle the components of striatal state dynamics that were 1) arousal-related, 2) task-related, and 3) person-specific. Arousal was indexed by the brain-wide mean systemic low frequency oscillation (msLFO) signal^28,40–42^.

This analysis revealed that striatal state dynamics were strongly linked with arousal, particularly during task-free, unconstrained resting-state scans (**Fig. 3**). Across the four resting-state and two Gambling task acquisitions, lower arousal (i.e., higher msLFO signal) was associated with more prevalent striatal coactivation bursts, both within (β_Arousal(Within)_ = 2.28, p < 0.001) and between subjects (β_Arousal(Between)_ = 0.31, p < 0.001), as well as with higher magnitude, more multi-input, and longer duration coactivation bursts.

**Figure 3.**
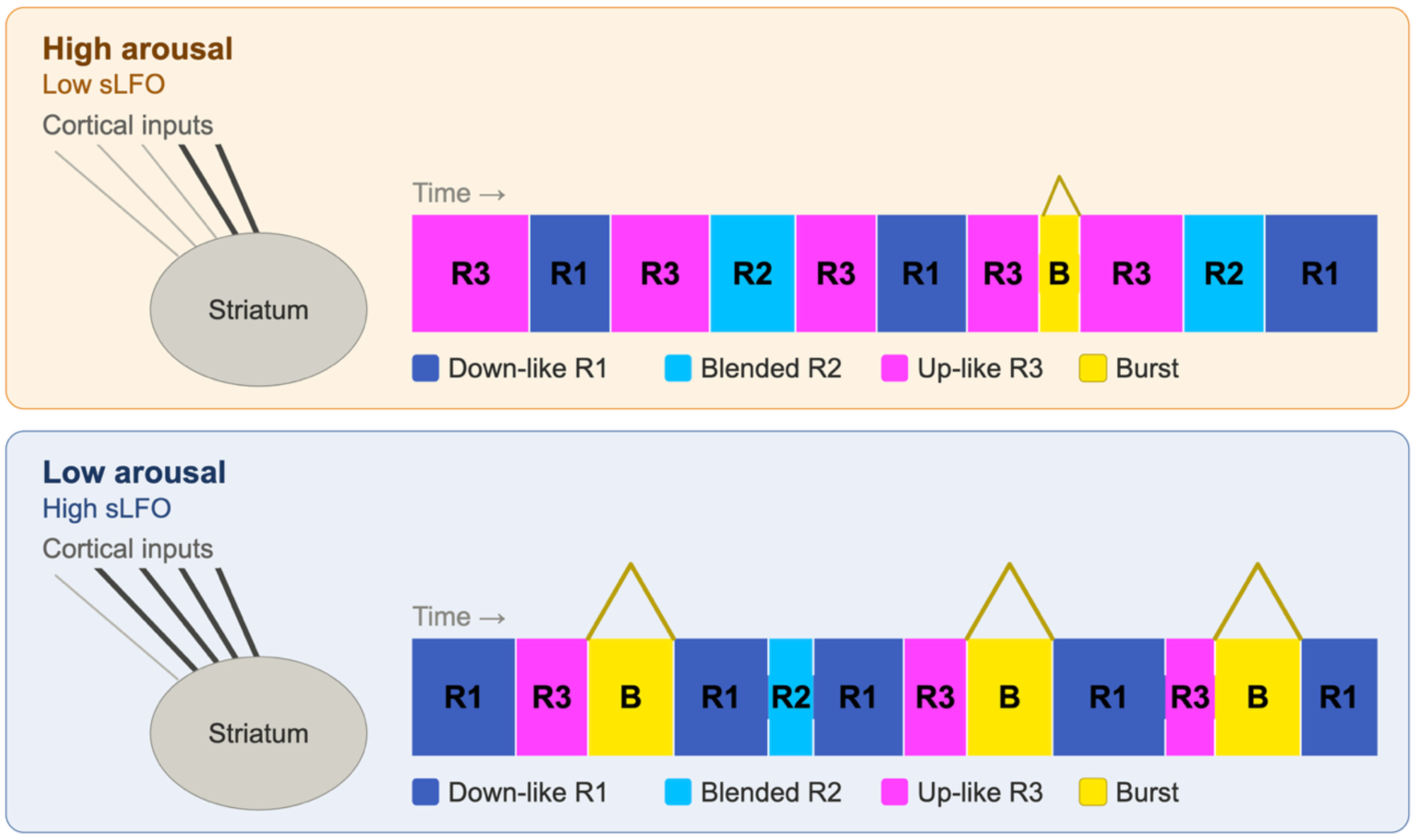
Striatal state dynamics during high and low arousal. Representative schematic depicting more prevalent and stable up-like rest 3 states during high arousal, versus more prevalent, stable, strong and multi-input coactivation burst states and more prevalent down-like rest 1 states during low arousal.

Task engagement attenuated arousal-linked bursting, β_Task x Arousal(Within)_ = −1.85, p = 0.007, while increasing task-related bursting, β_Task_ = 0.34, p < 0.001. Lower arousal was also associated with increased down-like rest 1 prevalence (β_Arousal(Within)_ = 1.93, p < 0.001) and duration (β_Arousal(Within)_ = 0.36, p < 0.001). Contrarily, lower arousal was associated with decreased blended rest 2 prevalence (β_Arousal(Within)_ = −3.27, p < 0.001; β_Arousal(Between)_ = −4.97, p < 0.001), and with decreased within-subject up-like rest 3 prevalence (β_Arousal(Within)_ = −1.07, p < 0.001). Up-like rest 3 states became significantly less stable during low arousal, displaying shorter dwell times (β_Arousal(Within)_ = −0.19, p < 0.001) and increased probability of transitioning out-of-state to burst state (β_Arousal(Within)_ = 0.01, p < 0.001) or down-like rest 1 (β_Arousal(Within)_ = 0.01, p < 0.001).

Overall, during low arousal, striatal state dynamics became more polarized (i.e., more burst and down-like rest 1 prevalence) while the stability of up-like rest 3 became degraded. During high arousal, bursting and down-like rest 1 states decreased, as up-like rest 3 states became more prevalent and stable.

### Functional Relevance of Striatal States During Task Engagement

In the absence of external demands at rest, coactivation burst state occupancy fluctuated around a stationary mean (**Fig. 4a**) in a manner linked to arousal level. Contrarily, during the Gambling task, coactivation burst state occupancy became time-locked to the task stimulation timeseries (**Figs. 4a-b, Supplementary Videos 3-4**). Between-subject analysis demonstrated that greater increases in striatum-wide burst state occupancy during win blocks relative to baseline were directly correlated with greater conventional striatum-wide BOLD activation during win blocks relative to baseline (i.e., contrast of parameter estimates; COPE), *t*(405)=2.66, p=0.008.

**Figure 4.**
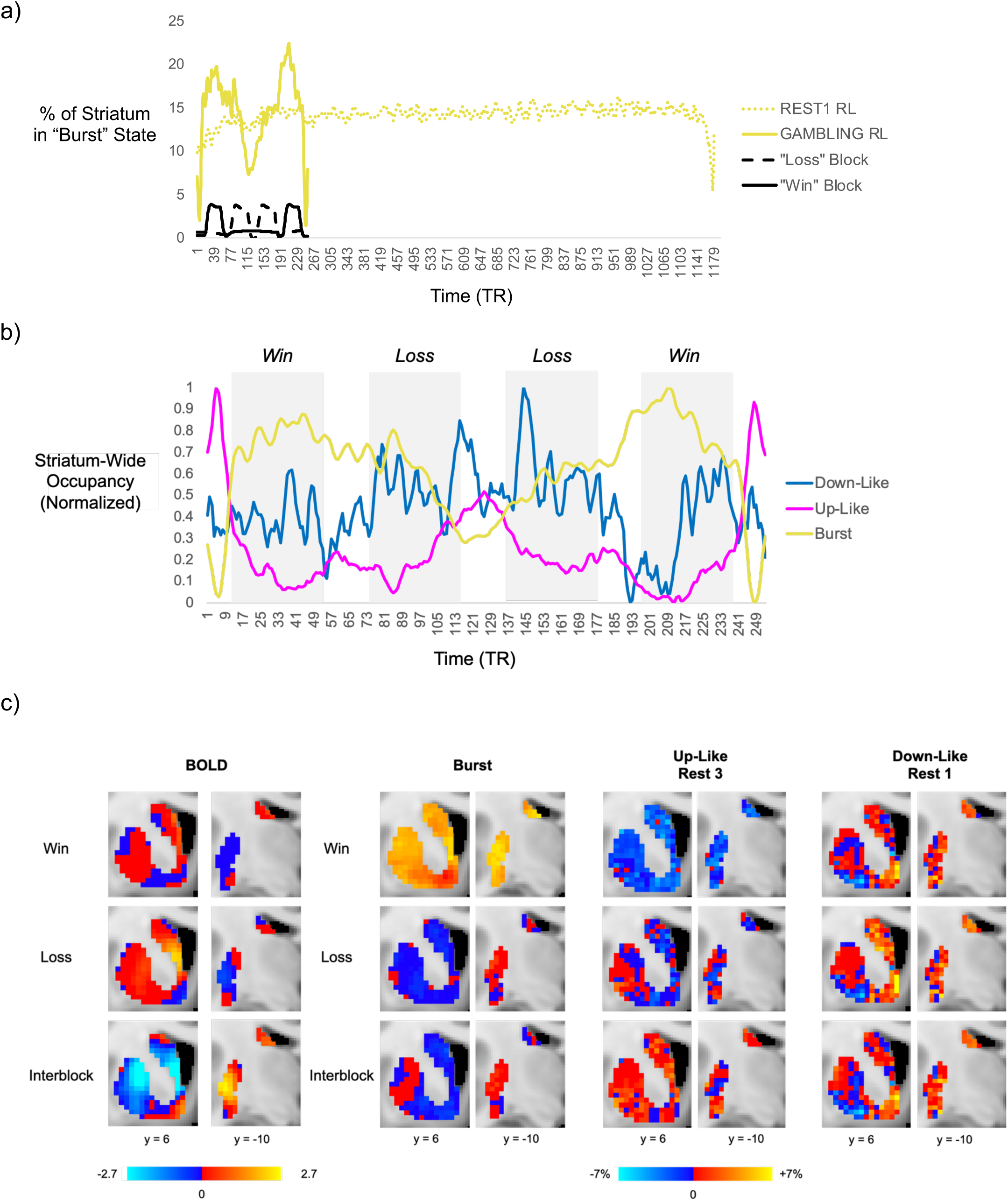
Burst coactivation state dynamics during the Gambling task. a) Group average time course of striatum-wide burst state occupancy during task versus resting-state. b) Group average time courses of striatum-wide state occupancy for burst, up-like and down-like states during the Gambling task across win blocks, loss blocks and inter-block intervals. c) Voxel-wise heat maps of (left) group average BOLD signal magnitude and (right) state occupancy relative to resting-state during different components of the Gambling task (win blocks, loss blocks, interblock intervals).

During inter-block intervals (i.e., the task baseline), coactivation burst occupancy was equivalent to that during resting-state scans (14.18% vs. 14.18%). However, burst-adjacent up-like rest state 3 occupancy during task inter-block intervals (33.54%) was significantly greater than that during resting-state scans (32.80%, *t*(406) = 4.32, *p* < 0.001), potentially reflecting a “task readiness” configuration. Against this inter-block baseline, win and loss blocks produced distinct reconfigurations of burst and rest coactivation states (**Fig. 4b**).

Relative to task inter-block intervals, win blocks significantly increased coactivation burst state occupancy (14.18%, vs. 17.95%, *t*(406) = 24.12, *p* < 0.001) and decreased up-like rest 3 occupancy (33.54% vs. 30.22%, *t*(406) = −16.66, *p* < 0.001), without altering down-like rest 1 occupancy (35.97% vs. 36.00**%**, *t*(406) = −0.19, *p* = 0.847) (**Fig. 4b**). In contrast, loss blocks increased down-like rest 1 occupancy (36.00% vs. 36.62%, *t*(406) = 4.03, *p* < 0.001) and decreased up-like rest 3 occupancy (33.54% vs. 32.20%, *t*(406) = −6.28, *p* < 0.001). Mean burst levels during loss blocks (14.30%) did not significantly differ from those during inter-block intervals (14.18%, *t*(406) = 0.50, *p* = 0.615), but certain loss blocks (i.e., those coming after a win block) elicited precipitous drops in burst levels to well below the resting-state average (**Fig. 4a**). Moreover, burst occupancy during loss blocks was significantly lower than that during win blocks (14.30% vs. 17.95%, *t*(406) = −15.70, *p* < 0.001).

Spatial comparison of group-average BOLD signal magnitudes (**Fig. 4c**, left panel) and coactivation state occupancies (**Fig. 4c**, right panels) during different Gambling task blocks illustrates how aggregate BOLD signal can both reflect and obscure underlying state-level dynamics. Consistent with prior analyses of this task and cohort^43,44^, ventral caudate was identified as a hotspot of BOLD signal increase – particularly during loss blocks – while nucleus accumbens showed appreciable BOLD signal increase only during inter-block periods, when reward anticipation is expected to be highest.

Win blocks elicited a striatum-wide increase in burst state occupancy, with peaks in ventral caudate and posterior dorsal putamen. This was accompanied by a striatum-wide decrease in up-like rest state 3 occupancy, consistent with reward-driven recruitment of burst states from the up-like pool. Notably, posterior dorsal putamen exhibited elevated burst state occupancy during win blocks despite low average BOLD signal, underscoring the capacity of time-resolved state dynamics to capture coordinated corticostriatal activity that is obscured in the aggregate BOLD signal. The nucleus accumbens, which exhibited minimal BOLD activation during win blocks, also displayed an increase in down-like rest state 1 occupancy during win blocks, indicating low coordination with cortex. Loss blocks produced a largely opposing spatial pattern: a striatum-wide increase in down-like rest state 1 occupancy with a corresponding reduction in up-like rest state 3 and burst occupancy. An exception was posterior dorsal putamen burst occupancy, which remained elevated throughout the task irrespective of block valence, potentially reflecting sustained motor response preparation throughout the task. Finally, during inter-block intervals, average BOLD signal was predominantly negative, yet up-like rest state 3 occupancy was selectively elevated without a corresponding increase in burst activity — a dissociation not recoverable from average BOLD alone — consistent with a task-readiness configuration in which the striatum maintains response readiness without escalating to suprathreshold output.

Overall, striatal coactivation state dynamics during the Gambling task appear to reflect a functionally relevant, multi-tiered system that flexibly reconfigures in response to task demands. Selective elevation of up-like rest 3 states during inter-block intervals may reflect a response readiness configuration that is mobilized during win blocks via transitions from up-like rest 3 to burst, and which is attenuated during loss blocks via transitions from up-like rest 3 to down-like rest 1.

### Individual Differences in Striatal State Dynamics and Behavior

Beyond the selective increase in up-like rest state 3 occupancy during inter-block intervals at the group level, individual-differences analyses provided convergent support for the interpretation of up-like rest 3 and down-like rest 1 as divergent task readiness configurations.

First, faster reaction times (RTs) during active blocks of the Gambling task were selectively associated with greater up-like rest state 3 occupancy relative to down-like rest state 1 occupancy during inter-block intervals, even after residualizing out msLFO-indexed arousal-related variance (e.g., overall median RT on “larger” trials: *t*(393) = −2.26, *p* = 0.024) (**Fig. 5a**). Conversely, slower RT was selectively associated with greater occupancy in the down-like rest 1 state during inter-block intervals (e.g., overall median RT on “larger” trials: *t*(393) = 2.80, *p* = 0.005) (**Fig. 5b**). These relationships held across almost all variations of Gambling task RT measures; regressions controlled for age and sex.

**Figure 5.**
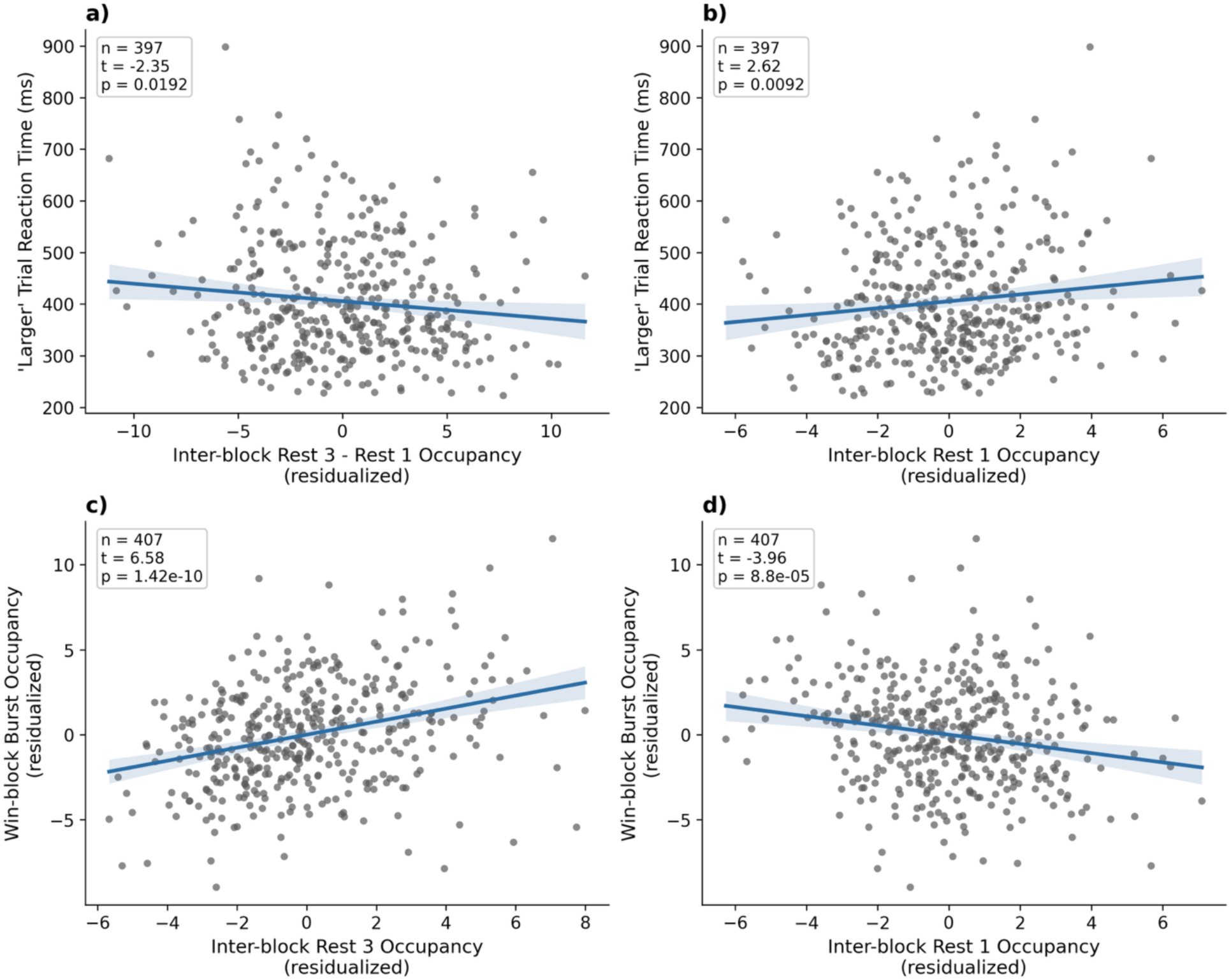
Individual differences in striatal state dynamics and Gambling task performance. Relationship between median reaction times (RTs) on trials of the Gambling task that received a “larger” prediction and striatum-wide inter-block interval occupancy of a) up-like rest 3 relative to down-like rest 1 and b) down-like rest 1. Relationship between win-block burst state occupancy (indexing reward reactivity) and c) inter-block up-like rest 3 occupancy and d) down-like rest 1 occupancy. Behavioral and neural metrics are averaged across both runs (i.e., RL and LR) of the Gambling task.

Second, greater reactivity to reward – indexed by burst coactivation occupancy during win trials – was also strongly associated with up-like rest 3 versus down-like rest 1 occupancy during inter-block intervals (**Fig. 5c-d**). Subjects with greater inter-block rest 3 occupancy showed greater win-block burst occupancy (*t*(403) = 6.58, p < 0.001). Conversely, subjects with greater inter-block down-like rest 1 occupancy showed reduced win-block burst occupancy (*t*(403) = −3.96, p < 0.001). Together, these findings highlight that behaviorally relevant signal is also carried by the low-coactivation rest states.

### Dynamic Reweighting of Frontal Cortical Coactivation Profiles

Finally, we demonstrate that the observed striatal state dynamics reflect functionally meaningful shifts in putative frontal input compositions. In individual subjects, we observed that burst states within a single striatal voxel could emerge from a diverse set of frontal coactivation configurations (**Fig. 6a**) - a reflection of striatal territories with fixed structural input profiles nonetheless exhibiting functional flexibility. The group-average profile underscored the functional relevance of these diverse configurations (**Fig. 6b**). For instance, we found significant, task block-dependent differences in frontal input composition during the prominent coactivation peaks visible after task-block onsets, as reflected in a significant block type × input interaction during this period, *F*(8, 18300) = 5.64, *p* < 0.001. Specifically, while ventral caudate coactivation with anterior insula increased significantly following both win block onset (*t*(406) = 6.25, *p* < 0.001) and loss block onset (*t*(406) = 3.12, *p* = 0.002) relative to inter-block onset, coactivation with OFC and vlPFC showed reward-selective early responses, with enhanced coactivation following win block onset relative to inter-block onset (*t*(406) = 3.33, *p* < 0.001; *t*(406) = 3.53, *p* < 0.001, respectively), but reduced coactivation following loss block onset relative to inter-block onset (*t*(406) = −3.41, *p* < 0.001; *t*(406) = −4.32, *p* < 0.001). Thus, beyond indexing aggregate increases or decreases in corticostriatal coupling, the observed striatal states also encoded meaningful changes in frontal input coactivation weightings. In the ventral caudate, this revealed putative salience-related insula engagement across win and loss blocks coupled with reward-selective engagement of OFC and vlPFC.

**Figure 6.**
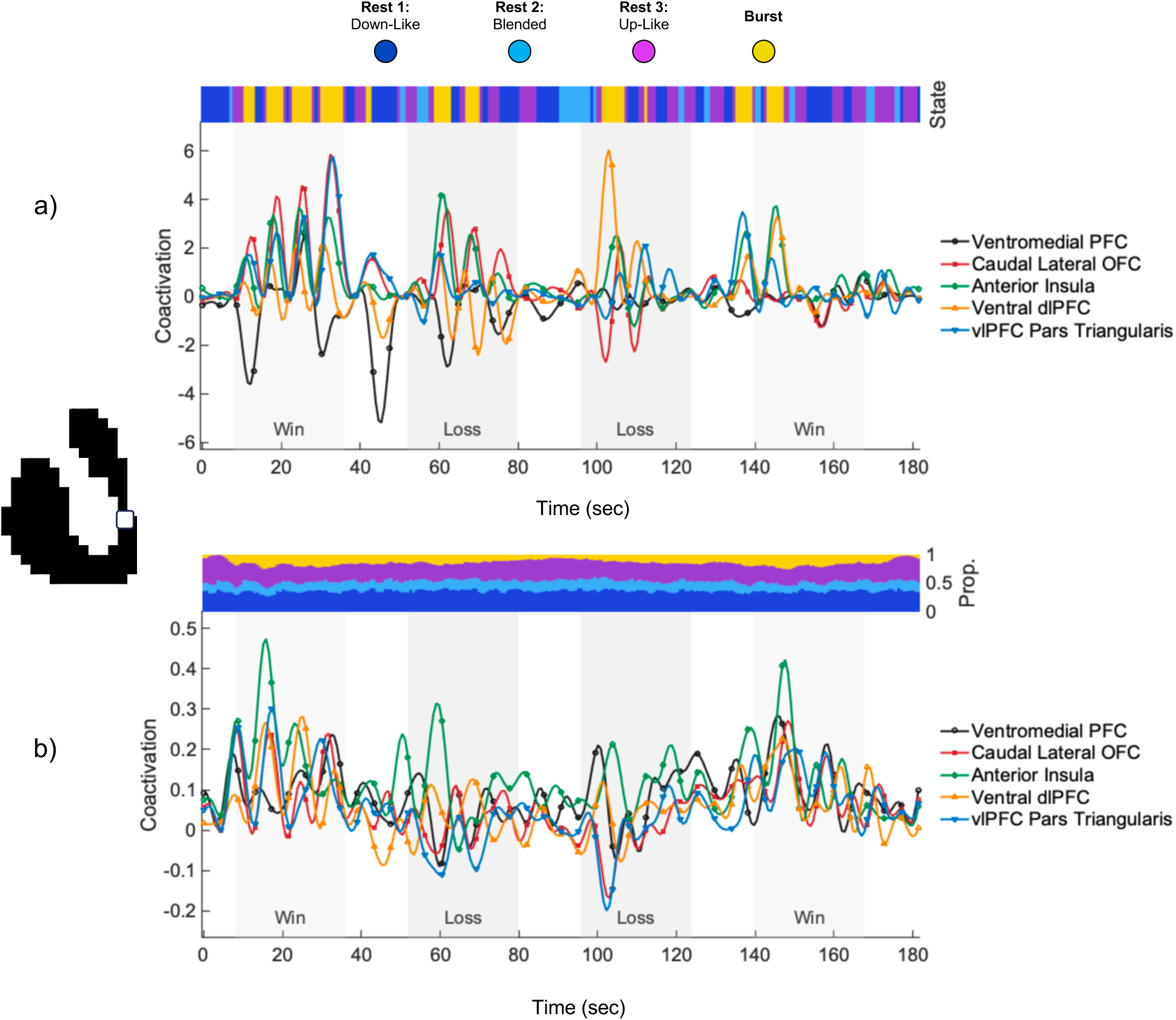
Dynamic reweighting of ventral caudate coactivation profile with frontal cortex during the Gambling task. a) Example individual-subject traces showing the time-varying coactivation profile of a ventral caudate voxel with its five dominant frontal cortical inputs across the Gambling task. Strip plot denotes the voxel’s coactivation state at each TR across the task. b) Group-average time-varying coactivation profile for the same ventral caudate voxel. Strip plot denotes the proportion of subjects whose ventral caudate voxel occupied each coactivation state at each TR.

## Discussion

This study leveraged neurobiologically informed circuit modeling and high temporal resolution computation of corticostriatal coactivation dynamics to identify and characterize mesoscale striatal state dynamics in human fMRI. Across more than 3 billion voxel-framewise striatal coactivation profiles with frontal cortical subregions, sampled over 65 minutes of scanning in 407 subjects, we identified four recurring states — down-like, blended, up-like, and burst — whose coactivation magnitudes, transition architecture, and task and arousal correlates show meaningful correspondence to canonical striatal electrophysiological states described in animal models. Importantly, these states reorganized systematically with both arousal and task context. At rest, burst dynamics were temporally irregular and strongly linked to arousal. During the Gambling task, by contrast, state occupancies became sharply structured by task epochs. Up-like rest state 3 increased during inter-block intervals in a manner consistent with a task-readiness configuration, and individual differences in inter-block state composition predicted reaction time and reward reactivity during active task blocks. Finally, the cortical composition of these states proved dynamically interpretable: the same ventral caudate territory expressed distinct frontal coactivation configurations across task contexts and subjects. Together, these findings suggest that human striatal function can be understood not only in terms of static corticostriatal connectivity architecture, but also in terms of transitions among a small set of recurring coactivation states that carry neurobiological and behavioral meaning.

### Correspondences and discrepancies with canonical striatal electrophysiology

The central result of this study is that striatal population coactivation states with frontal cortex, whose defining features recapitulate the logic of SPN states long characterized in animal models^21^, are discernible in human fMRI. As in the animal electrophysiology literature^21^, we observed two dominant rest-like states distinguished by low versus moderate corticostriatal coactivation, with strong bidirectional coupling between them and a marked asymmetry in their propensity to transition into a sparse, high-coactivation burst state. The burst state emerged preferentially from the up-like rest state, consistent with the principle that striatal firing requires prior depolarization into an up state before additional excitatory drive can trigger an output^45^. The behavioral and task-linked structure of these dynamics further supports the view that the identified states are not merely phenomenological, but mesoscale manifestations of a neurobiologically and behaviorally meaningful gating architecture.

The null-model results sharpen this interpretation. The empirical coactivation distribution departed from unimodality substantially more strongly than expected from autocorrelated noise, and the empirical burst regime was markedly more selective than the high-amplitude class identified when the same modeling procedure was applied to null data. These findings suggest that the observed states reflect meaningful characteristics of corticostriatal coactivation above and beyond the general statistical features of the data.

Several findings reflect limitations of the spatiotemporal blurring and indirectness of fMRI for recording striatal activity compared to invasive electrophysiology. First, the presence of the “blended” rest state 2. Electrophysiology data do not show a stable resting membrane potential for SPNs between the hyperpolarized down state and depolarized up state^21,45^. As such, the observation of a third, intermediary state in the fMRI data may reflect spatial pooling across populations of down and up state SPNs. Second, the coactivation profiles in each state display relatively large variance in average coactivation magnitude, and as such, the coactivation magnitude distributions of each state overlap to an appreciable extent. Electrophysiology data display smaller within-state variance in SPN membrane potential and clearer separation between states^21^. Third, the striatal state dwell times observed in the fMRI data (on the order of 3-5 seconds) are significantly longer than those observed in the electrophysiology data (i.e., 300-400 milliseconds for up and down states), reflecting the hemodynamic blurring inherent to fMRI and preprocessing-related smoothing. These discrepancies underscore that the current work is characterizing emergent properties of underlying cellular states at a mesoscale comprising seconds-worth of neuronal activity averaged across several hundred thousand neurons.

### Arousal as a tonic modulator of striatal state dynamics

A principal finding of the study is that striatal state dynamics were strongly coupled to arousal. During resting-state scans, lower arousal was associated with more frequent, higher magnitude, more multi-input, and longer duration coactivation bursts. This pattern is broadly consistent with prior work linking spontaneous high-amplitude coactivation events in resting-state fMRI to low-vigilance or drowsy states^28,46,47^. The present findings extend this literature by also demonstrating arousal-linked modulation of the sub-burst, low-coactivation rest state architecture. Lower arousal was associated with greater prevalence of the down-like rest 1 state and reduced stability of the up-like rest 3 state, suggesting that sustained maintenance of the task-ready, burst-adjacent rest 3 state may depend on adequate tonic arousal.

The mixed-effects decomposition of arousal-linked and task-linked contributions to state dynamics revealed that task engagement did not simply add more bursting on top of an arousal-determined baseline. The inter-block task baseline was characterized by elevated up-like rest state 3 without a net increase in burst states, and reward-evoked bursting appeared to reflect structured recruitment of that state rather than indiscriminate amplification of spontaneous burst propensity. This dissociation suggests that corticostriatal coactivation bursts operate in at least two partially separable modes: a tonic, arousal-linked mode at rest, and a phasic, task-linked mode during behavior. We demonstrate a principled means to separate burst-like activity that is behaviorally organized from endogenous burst-like activity that reflects arousal.

### Structured reconfigurations of striatal state dynamics during task engagement

The sharp task-locking and structured reconfigurations of striatal state dynamics during the Gambling task provide convergent evidence for their functional relevance. Relative to resting-state, task inter-block intervals showed increased up-like rest state 3 occupancy without increased bursting, suggesting a heightened-responsivity or task-readiness configuration. Against this inter-block baseline, win and loss blocks produced distinct state reconfigurations. Win blocks increased burst occupancy and reduced up-like rest 3 occupancy, consistent with recruitment of burst states from the up-like pool. Loss blocks, by contrast, increased down-like rest 1 occupancy and reduced up-like rest 3 occupancy, with mean burst occupancy remaining near inter-block baseline despite marked suppression during some loss epochs. This pattern supports a two-stage model in which the striatum first enters a task-ready, burst-adjacent configuration and is then either mobilized into burst output or relaxed into a less responsive configuration depending on behavioral context.

Importantly, these task-linked effects were not captured fully by aggregate BOLD signal. Inter-block periods, for example, were characterized by selectively elevated rest 3 occupancy despite predominantly negative average BOLD signal, whereas posterior dorsal putamen showed elevated burst occupancy during win blocks despite relatively low average BOLD. These dissociations highlight that time-resolved state dynamics can reveal organized corticostriatal coordination that is obscured when activity is averaged across time. In this sense, the current framework appears to recover latent structure that is only weakly visible in conventional mean-amplitude analyses.

The preferential localization of task-linked burst coactivation to ventral caudate further supports the functional plausibility of the framework. Ventral caudate sits within circuitry implicated in value integration, motivated choice, and action selection^48,49^. That these regionally specific dynamics emerged from a method that defines state structure with respect to each voxel’s own dominant frontal inputs, rather than an imposed anatomical parcel, argues that the approach is sensitive to meaningful functional topography rather than arbitrary regional averaging.

### Individual differences in striatal state dynamics and task performance

The individual-differences analyses provide additional support for the interpretation of up-like rest state 3 and down-like rest state 1 as divergent task-readiness configurations. Faster reaction times and greater reward reactivity during the Gambling task were associated with greater inter-block up-like rest 3 occupancy relative to down-like rest 1 occupancy, whereas slower reaction times and lower reward reactivity were associated with greater inter-block down-like rest 1 occupancy. In other words, subjects whose striatum more predominantly occupied the up-like relative to the down-like rest state during inter-block periods responded more quickly once the active task blocks began, and responded more vigorously in response to reward. This supports the idea that up-like rest 3 indexes a functionally relevant preparatory configuration rather than merely a statistical intermediary to burst coactivation. Notably, these effects persisted even after residualizing out msLFO-indexed arousal, suggesting that the relevant signal was not reducible to arousal-related fluctuation alone. Overall, these findings imply that subthreshold or non-burst state structure carries behaviorally meaningful information and that the balance between low-coactivation states may be as functionally important as the presence of overt burst coactivation events.

### Dynamic reweighting of frontal input composition

A further contribution of the present study is to show that the identified striatal states are interpretable not only in terms of overall coactivation magnitude, but also in terms of the putative contributions of distinct frontal inputs. Individual subject data demonstrated that burst coactivation states for individual striatal voxels could emerge from multiple different frontal coactivation configurations. Moreover, the group level data revealed that the observed state transitions are not only scalar changes in aggregate corticostriatal coupling, but also reflect recurring, functionally relevant changes in which frontal systems are relatively more or less engaged with a given striatal territory at a particular moment. Together, these data show how striatal territories with stable anatomical input architecture may nonetheless support flexible, context-sensitive computation^7^.

### Methodological considerations and limitations

While the identified states appear to be statistically and functionally meaningful, caution is warranted in interpretating them as sharply separable physiological categories. Their coactivation magnitude distributions overlap, measurement of their dwell times are blurred by hemodynamic smoothing, and one of the states—rest 2—likely reflects spatially mixed population configurations rather than a discrete electrophysiological regime. Moreover, although the null-model results strengthen support that the empirical burst regime is not a trivial consequence of autocorrelation or mixture fitting, the present study does not provide direct cellular validation of the identified states. In addition, reliability analyses indicated that single-run state-dynamic estimates are only modestly stable, whereas multi-run averages are substantially more reliable.

This supports the use of aggregated runs for individual-difference analyses – as done here – and cautions against overinterpreting single-run estimates in translational settings. Furthermore, the generalizability of the identified striatal state structure across brain hemispheres, acquisition protocols, developmental stages, and clinical populations remains to be explored. Finally, the correspondence between fMRI-based states and canonical SPN dynamics motivates direct validation against invasive electrophysiology, concurrent intracranial recordings, or other modalities capable of resolving faster neuronal timescales in future work.

### Conclusion

We identify and characterize striatal state dynamics in human fMRI that recapitulate key features of canonical electrophysiological SPN states and reorganize systematically with arousal, task context, and behavior. These findings show that neurobiologically meaningful corticostriatal dynamics can be recovered from high-quality fMRI data when analyzed at the level of framewise coactivation profiles. More broadly, they suggest that human striatal function can be understood not only in terms of static connectivity architecture, but also in terms of transitions among a small set of recurring population states. This provides a foundation for probing how corticostriatal dynamics support human behavior and how they may become dysregulated in neuropsychiatric disease.

## Methods

### Subjects

Data used in these analyses include fMRI scans from individuals gathered as part of the Human Connectome Project Young Adult (HCP-YA) 1200 subject release. Subjects consented to participate under the approval of the Washington University in St Louis Institutional Review Board. A detailed description of the recruitment for HCP-YA is provided by others^50–52^. Briefly, individuals were excluded by HCP-YA if they reported a history of major psychiatric disorder, neurological disorder, or medical disorder known to influence brain function. We further excluded individuals from the HCP-YA database that reported a family history of schizophrenia, met DSM-IV criteria for alcohol dependence, or reported a lifetime history of repeated substance use (>10 instances of cocaine, hallucinogen, opiate, sedatives, or stimulant use, >20 instances of tobacco use or >100 instances of marijuana use). In addition, participants included in the present analyses provided a breath sample indicating <0.05 blood alcohol content on the day of scan and a urine sample that was negative for any substances of abuse (cocaine, marijuana, opiates, amphetamine, or methamphetamine). Among 458 eligible individuals, a final sample of n=407 (Female = 281) had complete fMRI data for all six acquisitions of interest (REST1 RL, REST1 LR, REST2 LR, REST2 RL, GAMBLING RL, GAMBLING LR) who were an average age of 28.66 years of age (±3.65, range 22–36) and reported completing an average of 15.34 years of education (±1.59, range 11–17). 30 of these subjects met DSM-IV criteria for Alcohol Abuse at some point during their lifetime. In the past 7 days, 181 of these subjects reported drinking alcohol on zero days, 81 reported drinking on one day, 63 on two days, 41 on three days, 24 on four days, 12 on five days, 3 on six days, and 2 reported drinking on seven days. 292 subjects reported never having used marijuana during their lifetime; 82 reported using marijuana between one and five times; 12 reported between six and ten times; 21 reported between 11 and 100 times. Three hundred and eight participants self-identified as White, 56 identified as Black or African American, 32 identified as Asian, Native Hawaiian, or Pacific Islander, five identified as multiracial, and six were unknown or not reported.

### fMRI Data Acquisition

HCP-YA neuroimaging data were acquired with a standard 32-channel head coil on a Siemens 3T Skyra modified to achieve a maximum gradient strength of 100 mT/m^50–52^. Gradient-echo EPI images were acquired with the following parameters: TR = 720 ms, TE = 33.1 ms, flip angle = 52°, FOV = 280 × 180 mm, Matrix = 140 × 90, Echo spacing = 0.58 ms, BW = 2,290 Hz/Px. Slice thickness was set to 2.0 mm, 72 slices, 2.0 mm isotropic voxels, with a multiband acceleration factor of 8. Resting-state data were acquired across two sessions (REST1, REST2), each with two phase-encoding directions (LR and RL), yielding four resting-state runs (~14.4 minutes each). Participants were instructed to lie still with their eyes open and fixated on a bright crosshair on a dark background. In addition, two ~3-minute incentive salience (“Gambling”) task runs (GAMBLING RL and GAMBLING LR) were acquired in the same session as REST1.

### Gambling Task

The Gambling task (i.e., incentive processing task) was adapted from Delgado and Fiez^53^. Subjects guessed the number on a mystery card in order to win or lose money. Card numbers ranged from 1–9, and subjects were told to indicate if they thought the mystery card number was more or less than 5 by pressing one of two buttons on the response box. Feedback consisted of the number on the card (generated by the program as a function of whether the trial was a win, loss or neutral trial) and either: 1) a green up arrow with “$1&rdquo\semicolon \; for win trials\comma \; 2\rpar a red down arrow next to &minus\semicolon \; $0.50 for loss trials; or 3) the number 5 and a gray double headed arrow for neutral trials. The task was presented in blocks of 8 trials that were either mostly win (6 win trials pseudo randomly interleaved with either 1 neutral and 1 loss trial, 2 neutral trials, or 2 loss trials) or mostly loss (6 loss trials interleaved with either 1 neutral and 1 win trial, 2 neutral trials, or 2 win trials). All participants were provided with money for completing the task, though it was a standard amount across subjects.

### fMRI Preprocessing

We began with HCP-YA minimally preprocessed fMRI data^54^. Briefly, this preprocessing pipeline removes spatial distortions, realigns volumes to compensate for subject motion, registers the echo planar functional data to the structural data, reduces the bias field, normalizes the 4D image to a global mean, and masks the data with a final FreeSurfer-generated brain mask^54^. We performed further denoising of each subject’s data, including spatial blurring with a 6-mm full-width half-maximum Gaussian kernel and temporal filtering (0.01<f <0.1 Hz). Next, the time series of each voxel was confound regressed in a GLM with 17 regressors of no interest: six motion parameters (three translations and three rotations) obtained from the rigid-body alignment of EPI volumes and their six temporal derivatives; the mean time series extracted from white matter; the mean times series extracted from CSF; and a second-order polynomial to model baseline signal and slow drift. Finally, we applied RIPTiDe using “rapidtide”, a performance optimized open source implementation of the RIPTiDe processing stream (https://github.com/bbfrederick/rapidtide), described in detail previously^55^. RIPTiDe removes the systemic low frequency oscillation (sLFO) signal from the data, which was recently found to introduce a strong temporally structured artifact into FC estimates in fMRI data^28^.

### Regions of Interest

The right striatum ROI comprised of 1710 voxels was defined using a mask from previous work^7,8^, which was generated by averaging the union of the caudate, putamen, and nucleus accumbens FreeSurfer parcellations of each of 77 low-head motion HCP-YA subjects^56^, intersected with a replication subsample and constrained to remove edge voxels. Frontal cortical regions of interest comprised the 21 right hemispheric frontal cortical parcels in the Schaefer 100-parcel, 7-network atlas^33^.

### Constructing Corticostriatal Functional Connectivity Profiles

For each subject, we computed static functional connectivity between each of the 1710 right striatal voxel and each of the 21 ipsilateral frontal cortical ROIs as the Pearson correlation between each pair’s z-scored BOLD time series. These voxel-by-ROI connectivity matrices were then averaged across all subjects to generate a group-average static connectivity map. For each striatal voxel, we identified the five frontal cortical ROIs with the highest group-average static functional connectivity and defined this set as the voxel’s connectivity profile. This approach assumes that corticostriatal temporal dynamics arise primarily from fluctuations in the strength of drive along a stable set of afferent pathways, rather than from rapid changes in which cortical regions are connected. Holding the top five inputs fixed per voxel thus provided an anatomically informed dimensionality reduction.

### Temporal Unwrapping to Yield Framewise Coactivation Profiles

Next, for each striatal voxel in each subject, we applied temporal unwrapping^27^ to each functional connection in the connectivity profile. For each pair of striatal voxel *v* and frontal cortical ROI *r*, we computed the element-wise product of their z-scored BOLD timeseries, yielded an “edge timeseries” encoding the pair’s coactivation magnitude (i.e., correlation product, CP) at each TR *t*:

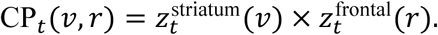

Mathematically, the time-average of CP_*t*_(*v*, *r*) across all TRs is equal to the Pearson correlation (i.e., static functional connectivity) between *v* and *r*. Thus, this procedure decomposes the static functional connectivity of each corticostriatal connection into a framewise measure of coactivation magnitude at the temporal resolution of the scan (TR = 0.72 s).

Within this edge-time-series framework, positive and negative CP values have a specific mathematical interpretation. Positive CP values occur when the striatal and cortical BOLD signals deviate from their respective means in the same direction at a given TR — that is, when both are relatively high or both are relatively low. Negative CP values occur when the two signals deviate in opposite directions, with one above and the other below its mean. Thus, positive CP values index momentary same-signed co-fluctuation, whereas negative CP values index momentary opposite-signed co-fluctuation. In the present corticostriatal application, we used these edge-time-series values as an fMRI-derived measure of momentary coordination between each striatal voxel and its strongest frontal cortical inputs.

Collectively, this analysis yielded, for each striatal voxel and TR, a five-dimensional vector quantifying the coactivation magnitude between the striatal voxel and each of its five strongest frontal cortical inputs. These “coactivation profiles” were stacked across all time points, striatal voxels, and subjects for a given scanning acquisition to form a matrix upon which subsequent clustering analysis was performed to identify recurring coactivation states. To facilitate comparisons across fMRI acquisitions (e.g., between resting-state and task), we established a common reference scale for coactivation magnitude by computing the mean *μ* and standard deviation *σ* of all coactivation magnitudes in one resting-state acquisition (REST1 LR).

### Clustering “Burst” and “Rest” Coactivation Frames

Next, we applied two-component Gaussian mixture modeling to the matrix of coactivation profiles to separate disproportionately high-coactivation “burst” states from lower-coactivation “rest” states. Before GMM fitting, we first rectified each frame-voxel’s 5-dimensional coactivation vector by setting negative values to zero. This choice was motivated by the neurobiological constraint that corticostriatal projections are excitatory, and electrophysiological evidence that convergent cortical excitation can summate to amplify striatal activity.

Accordingly, positive coactivation values were allowed to accumulate across cortical inputs, whereas negative coactivation with one input was not treated as mechanistically subtracting from positive coactivation with other inputs. We then computed the mean of the five rectified inputs (“mean-of-5” amplitude) for each frame-voxel as a summary of overall positive coactivation strength.

We then fit two-component Gaussian mixture models to (i) the pooled distribution of rectified coactivation values across all inputs and frame–voxels, and (ii) the pooled distribution of mean-of-5 amplitudes. For each distribution, we defined a threshold as the positive decision boundary separating the lower-mean and higher-mean Gaussian components. This yielded 1) a set of per-input thresholds *τ*_*j*_ indicating the magnitude above which input *j* is considered to have disproportionately high coactivation with a striatal voxel, and 2) a mean-of-5 threshold *τ*_mean_ indicating when the combined positive coactivation strength across all inputs is considered to be disproportionately high. These thresholds were estimated from one resting-state acquisition (REST1 LR) and then held fixed across all other resting-state and task acquisitions, providing reference-standardized thresholds.

A frame–voxel was labeled as a coactivation burst if two criteria were met: (1) its mean-of-5 amplitude exceeded *τ*_mean_, and (2) at least one of its five inputs exceeded its per-input threshold (a 1-of-5 coordination rule). This hybrid criterion ensured that (i) overall coactivation magnitude was disproportionately high and (ii) coactivation magnitude with at least one individual input was disproportionately high. For each subject, we summarized coactivation burst dynamics by calculating: the fraction of all frame–voxels classified as bursts, the mean amplitude of positive inputs on burst frames, and the distribution of bursts as a function of how many of the five inputs were supra-threshold (k-of-5 composition).

We next characterized the remaining background organization of corticostriatal coactivity. We began by restricted the frame–voxel coactivation matrix to rows corresponding to non-burst frames and randomly subsampling 200,000 rows to use as a training set. Using their 5-dimensional coactivation vectors, we computed a frame-by-frame similarity matrix quantifying how similar the coactivation pattern is between any two non-burst frame–voxels. We then applied unsupervised Louvain community detection^37,38^ to this similarity matrix to partition the training frames into *K*_rest_communities (rest coactivation states) that maximize within-state similarity and minimize between-state similarity.

For each state, we computed its centroid, defined as the mean 5-dimensional coactivation vector across all training frames assigned to that state. To obtain a consistent ordering of states across runs, we sorted state labels by their mean signed coactivation strength, such that low-index states correspond to lower overall corticostriatal drive and higher-index states to higher drive. These centroids define a low-dimensional basis of canonical resting coactivation patterns. Every non-burst frame–voxel in the full dataset (training and held-out) was then assigned to the rest state whose centroid had the highest cosine similarity with its 5-dimensional coactivation vector.

### Null Modeling

We compared the empirical coactivation magnitude distribution against a null distribution generated from autoregressive model of order one (AR(1)) surrogate datasets. For each subject, independent surrogate timeseries were generated separately for each striatal voxel and frontal cortical ROI by estimating the lag-1 autocorrelation coefficient from the corresponding preprocessed BOLD signal and simulating a new timeseries with matched mean, variance, and autocorrelation structure. These surrogate timeseries therefore preserved the first-order temporal autocorrelation and marginal distributional scale of each regional signal, while disrupting coordinated corticostriatal co-fluctuation structure. For each of 1000 null iterations, the full coactivation computation pipeline was reapplied to the surrogate data, including z-scoring, computation of framewise corticostriatal correlation products, selection of each striatal voxel’s five strongest frontal cortical inputs, rectification of negative coactivation values, common-scale normalization. For each iteration, we fit both a one-component and two-component GMM to the distribution and computed the BIC for each model and also computed the percent of frame-voxels classified as coactivation bursts under the two-component model. This allowed for comparison with the empirical data.

### State occupancy, dwell time, and transitions

In the empirical data, we derived several metrics to characterize the corticostriatal coactivation state dynamics:

*Occupancy.* For each subject and state, we computed the proportion of all frame–voxels assigned to that state, yielding subject-level occupancy fractions for each coactivation rest state and for coactivation bursts. We also computed global occupancy across all subjects. These measures index how much of a subject’s striatal fMRI data is spent in each coactivation state.

*Dwell times.* For each voxel, state, and subject, we identified contiguous runs of TRs in which the voxel remained in the same state. We defined dwell time as the number of consecutive TRs within a run and summarized, for each state, the average dwell time per subject and the group mean and standard error. This assesses how long striatal areas tend to remain in each coactivation state before transitioning.

*State transitions.* To characterize transitions between states, we examined successive TR pairs within each voxel’s time series and tallied all instances in which state *i* at TR *t* was followed by state *j* at TR *t* + 1. For each subject, we normalized the resulting state-by-state transition count matrix by row to obtain a transition probability matrix and then averaged these across subjects.

*Burst composition across inputs.* Finally, to better understand coactivation bursts from the cortical context, we examined, on coactivation burst frames only, which subset of a voxel’s five inputs exceeded their per-input thresholds. We encoded each pattern of supra-threshold inputs as a 5-bit mask and computed, for each voxel, the frequency and percentage of coactivation burst frames associated with each input combination. Aggregating across subjects produces voxelwise “coactivation burst motif” profiles that describe which combinations of canonical frontal inputs most commonly drive coactivation bursts in that voxel.

### Linear Mixed-Effects Modeling

Using run-level msLFO amplitude as a physiological arousal index, we fit linear mixed-effects models across all six runs per subject to decompose each burst metric:

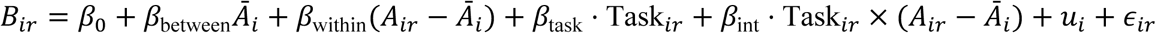

where *Ā*_*i*_ is the subject’s mean msLFO across all six runs, (*A*_*ir*_ − *Ā*_*i*_) is the within-person run-specific deviation, Task is 0 for REST and 1 for GAMBLING, and *u*_*i*_ is a subject random intercept. The person-specific stable component was defined as each subject’s mean model residual across all six runs. The task-effect residual was defined as the subject’s mean residual on task runs minus mean residual on rest runs.

### Behavioral Regression Analyses

For each behavioral outcome and each neural predictor, we fit:

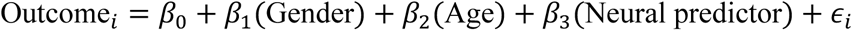

where the neural predictors were subjects’ mean residuals on task run minus mean residuals on rest runs from the linear mixed-effects decomposition, in order to remove arousal effects.

## Data Availability

The HCP-YA dataset is publicly available on an open access repository (https://db.humanconnectome.org/app/template/Login.vm), which can be accessed after signing a data use agreement.

The striatal and frontal cortical masks used for analysis are freely available for download at: https://neurovault.org/collections/21499/.

## Code Availability

All custom code for identification and analysis of striatal states are available at: https://github.com/ckorponay/Corticostriatal-Temporal-Dynamics.

