## Supplemental Figures for "Human striatal population state dynamics"

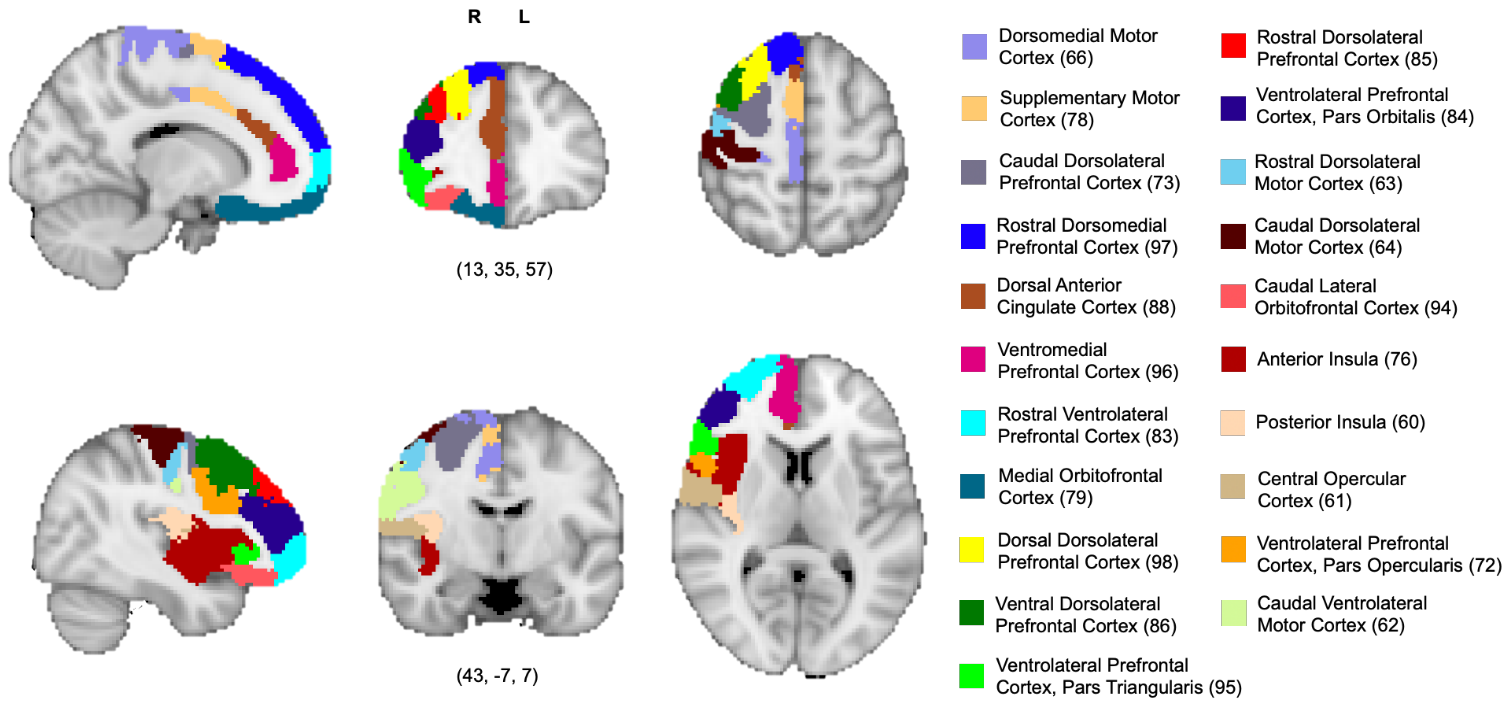


**Figure S1.** *Frontal cortical regions of interest.* The 21 right hemispheric frontal cortical ROIs in the Schaefer 100-parcel, 7-network atlas. Numbers refer to the ROI label in the atlas.

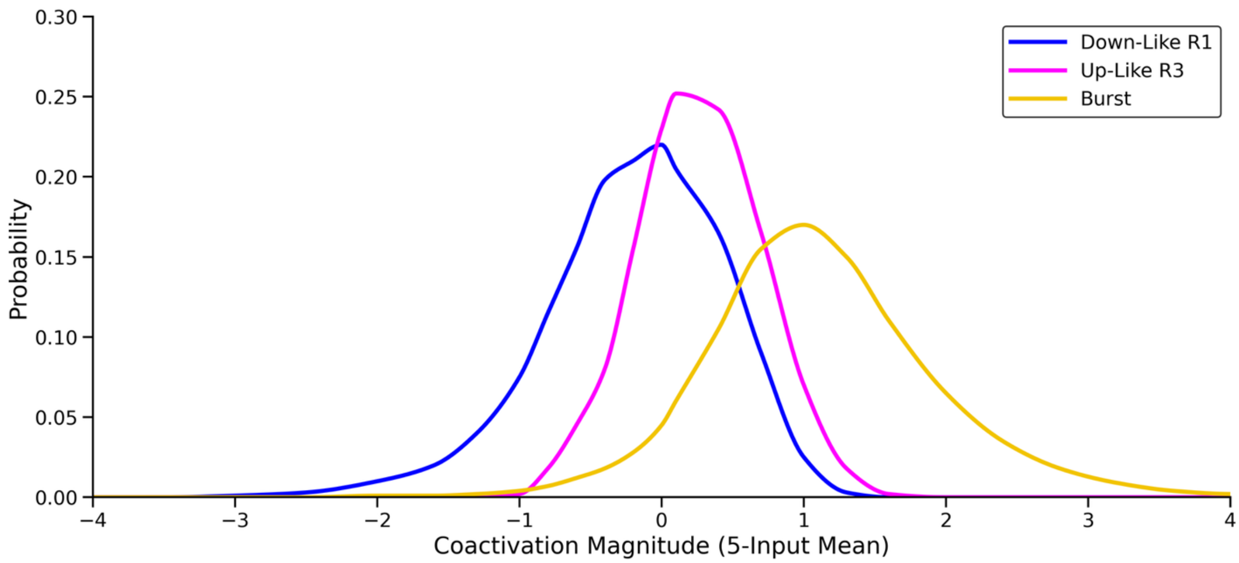


**Figure S2.** *Distribution of mean coactivation magnitudes by state.* For voxel-frame-wise striatal coactivation profiles classified to each state, the frequency of various coactivation magnitudes across the four resting-state acquisitions.
